# Statistical profiling reveals correlations between the cell response to and structural sequence of Rho-GAPs

**DOI:** 10.1101/2020.09.20.305599

**Authors:** Na Kang, Tsubasa S. Matsui, Shinji Deguchi

**Affiliations:** Division of Bioengineering, Graduate School of Engineering Science, Osaka University, Toyonaka, Japan

**Keywords:** Rho family of GTPases, Rho-GTPase-activating proteins, epithelial-mesenchymal transition, morphology, migration, statistical analysis

## Abstract

Rho-GTPase-activating proteins (Rho-GAPs) are essential upstream regulators of the Rho family of GTPases. Currently, it remains unclear if the phenotypic change caused by perturbations to a Rho-GAP is predictable from its structural sequence. Here we analyze the relationship between the morphological response of cells to the silencing of Rho-GAPs and their primary structure. For all possible pairs of 57 different Rho-GAPs expressed in MCF10A epithelial cells, the similarity in the Rho-GAP silencing-induced morphological change was quantified and compared to the similarity in the primary structure of the corresponding pairs. We found a distinct correlation between the morphological and structural similarities in a specific group of RhoA-targeting Rho-GAPs. Thus, the family-wide analysis revealed a common feature shared by the specific Rho-GAPs.

## 1. Introduction

The Rho family of GTPases (Rho-family), with three members of Cdc42, Rac1, and RhoA, is a master regulator of the actin cytoskeleton. As its upstream regulators, in humans there are 66 different Rho-GTPase-activating proteins (Rho-GAPs) known to negatively control Rho-family by inactivating its GTPase activity [1-3]. The Rho-GAP domain is a specific domain commonly found in the structure of Rho-GAPs and responsible for the Rho-GTPase activity. They also contain FBAR, CRAL-TRIO, and Ras-associating domains, which are known to jointly mediate membrane remodeling pathways, small lipophilic molecule binding, and cancer progression [4-7]. The presence of such large numbers and varieties of Rho-GAPs is supposed to allow the cells to exhibit a wide range of functions using the actin cytoskeleton in a cell context-dependent manner. Indeed, Rho-GAPs have been implicated in diverse processes such as migration, proliferation, morphogenesis, and tumorigenesis [8-10]. In this regard, specific Rho-GAPs have often been sporadically studied, but attempts have recently begun to more comprehensively understand the role of Rho-GAPs. For example, molecular structural and functional similarities [11] and subcellular localizations and associated molecules [12] were investigated at the Rho-GAP family-wide level.

We also recently investigated the effect of silencing individual Rho-GAPs on the morphology of epithelial cells with an aim of identifying critical factors associated with the epithelial-mesenchymal transition (EMT) [13]. For this experiment, we conducted a two-step screening for human mammary epithelial MCF10A cells cultured at confluent (1st screening) and sparse (2nd screening) conditions, respectively, and consequently 15 (specifically, ARHGAP4, ARHGAP21, ARHGAP11A, ARHGAP17, DEPDC1B, ABR, BCR, MYO9B, ARAP2, PIK3R1, OCRL, GMIP, SH3BP1, SRGAP1, and SRGAP2) and 2 (ARHGAP4 and SH3BP1) Rho-GAPs were found -among 57 different ones expressed in the cells - to play a significant role in maintaining the original cell shape in confluent and sparse cultures, respectively. However, our focus in this study was primarily on the role and activation mechanism of the most effective one, ARHGAP4. Thus, the family-wide nature of Rho-GAPs in the maintenance of cell morphology remains to be explored.

Here we provide in-depth analysis to find if there is a specific relationship between the sequential information of Rho-GAPs and resulting phenotypes expressed in MCF10A cells (Fig. 1). The phenotypes we examined were the cell morphology and migratory potential, the latter of which is another factor to be regulated in EMT and associated with the activity of Rho-GAPs [14, 15]. Through the analysis, we revealed the presence of a correlative relationship shared by specific Rho-GAPs that target only RhoA but not Cdc42 and Rac1, which suggests that the more similar their primary structures are, the more similar morphological change cells undergo upon the silencing of the Rho-GAPs. We also found a cell density-dependent relationship, which correlates the cell shape change due to the silencing of specific Rho-GAPs with the migratory potential. These relationships provide a basis for predicting the cell response upon perturbations to the Rho-GAPs and for considering how phenotypes are dictated in the presence of the variety of Rho-GAPs.

**Fig. 1.**
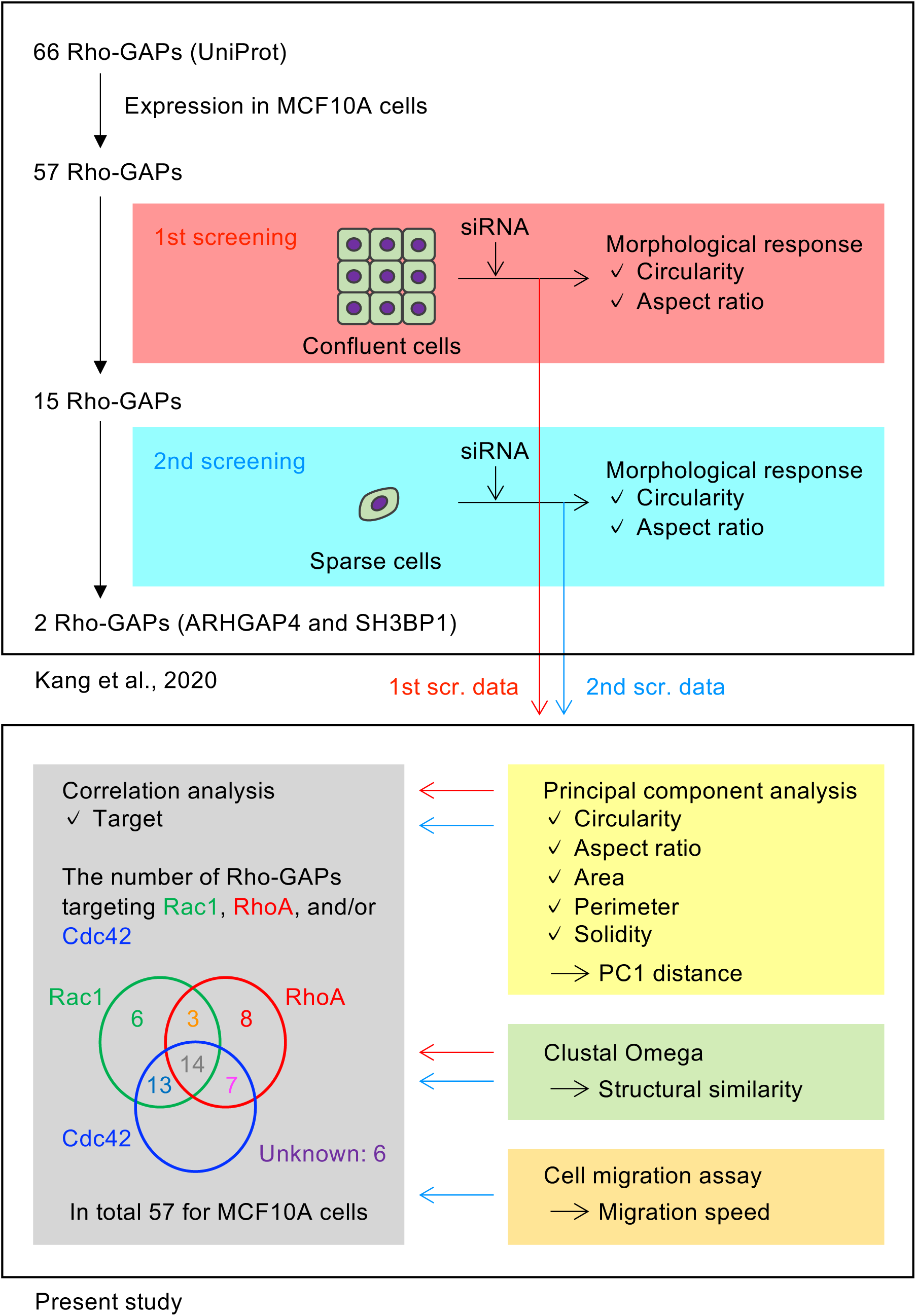
Schematic illustration of the present analysis (below) and the relationship with the previous experiments (Kang et al., 2020) [13] (upper).

## 2. Results

### 2.1. Comprehensive visualization of how silencing of individual Rho-GAPs affects the morphology of MCF10A cells

For 1st screening in the previous study [13], MCF10A cells at confluent conditions were subjected to siRNA-mediated silencing of each of all the 57 Rho-GAPs (Supplementary Fig. S1, S2 for representative images and methodologies). To capture the resulting features of cell morphology, principal component analysis (PCA) was performed on 5 parameters, Area, Perimeter, Aspect ratio (AR), Circularity, and Solidity. The first (PC1) and second (PC2) principal components reached a contribution of 57.6% and 37.6% of the explained variance, respectively (Fig 2A). PC1 is predominantly determined by the combination of AR, Circularity, and Solidity, while PC2 is approximately determined by Area and Perimeter. Circularity and Solidity were found to exhibit a similar response in confluent MCF10A cells that possess relatively smooth outlines with no marked concave morphology (Supplementary Fig. S1, S2). Consequently, a higher value of PC1 is interpreted to display a rounder cell shape, while that of PC2 represents a cell that is bigger in size. The relative importance of AR and Circularity in PC1 over the other 3 parameters does justify our previous choice [13], in which only the former two were adopted to detect EMT. Our PCA result showed that the 15 Rho-GAPs -selected in 1st screening -were all located at the left half-plane of the PC1-PC2 coordinate with an increased AR and a decreased Circularity compared to control. This tendency seems reasonable in that in general epithelial cells lose a round shape upon EMT to become elongated [16, 17]. Here note that, to identify the reliability of the perturbation, the siRNA transfection effectiveness of these 15 Rho-GAPs has been affirmed at the mRNA level [13]. Hierarchical cluster analysis (HCA) was performed based on the PC1 data to show a dendrogram that visualizes the clustering (Fig. 2B). Indeed, all the selected 15 Rho-GAPs were located in nearby clusters (7 in red and 8 in blue).

**Fig. 2.**
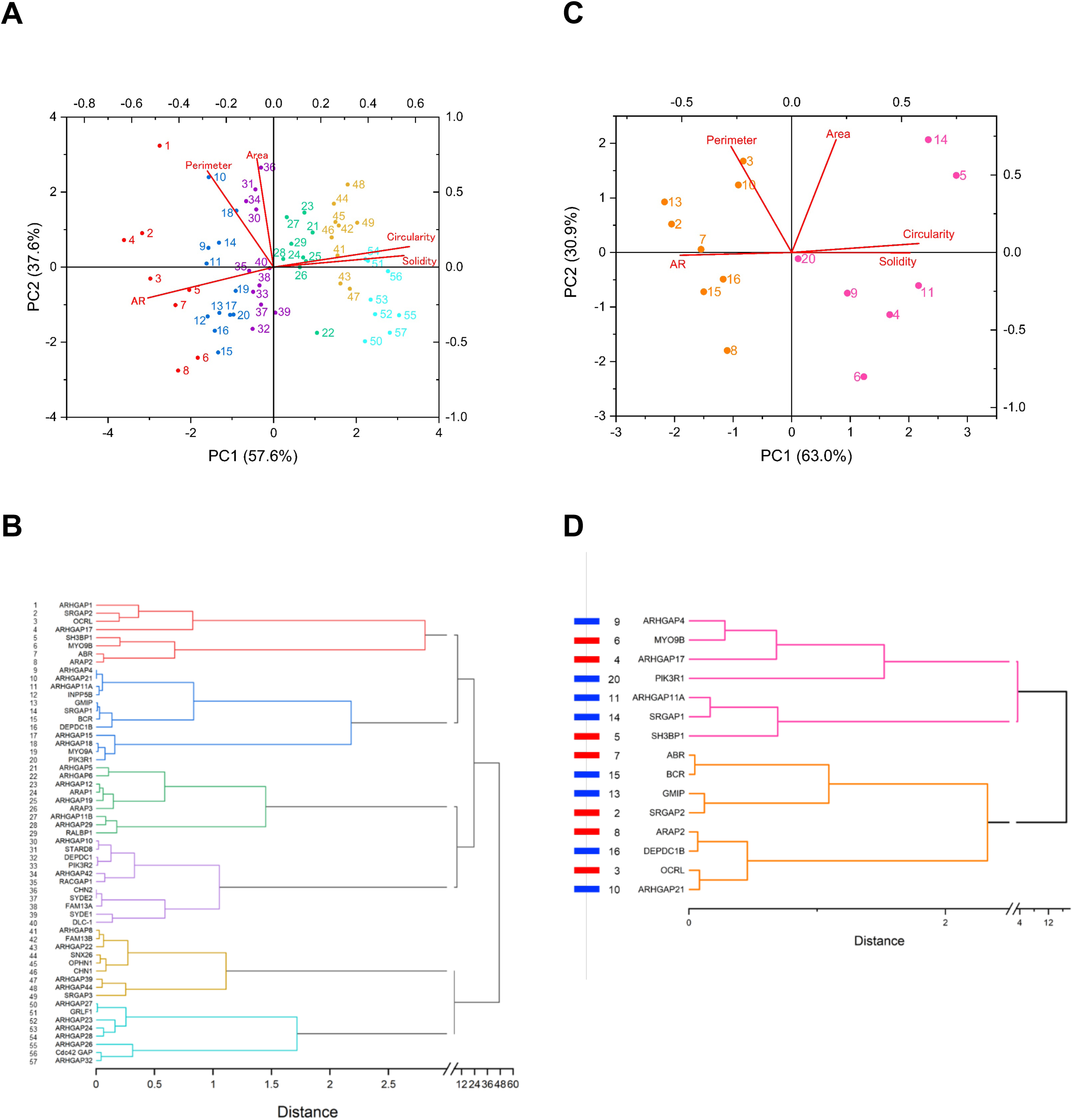
PCA and HCA visualize the morphological response of MCF10A cells to the silencing of Rho-GAPs. (A) PCA for confluent cell data. The numbers indicate each of the 57 Rho-GAPs, the names of which are specified in B. See Supplementary Table S1 for more information. (B) HCA based on the PC1 data of A. 6 major clusters, shown by different colors (consistent with those in A), were obtained. (C) PCA for sparse cell data. The numbers indicate each of the 15 Rho-GAPs, which are specified in D. See Supplementary Table S2 for more information. (D) HCA based on the PC1 data of C. 2 major clusters, shown by different colors (consistent with those in C), were obtained.

We originally intended to search for Rho-GAPs involved in the maintenance of cell morphology independent of cell density. To do this, we narrowed down the range of Rho-GAPs from 1st screening data examined in confluent culture to find, in 2nd screening, ones that are effective as well in sparse culture (Supplementary Fig. S1). From the PCA on the 15 Rho-GAPs that passed 1st screening, we found that PC1 for these cell populations is also approximately determined by AR, Circularity, and Solidity with a contribution of 63.0% of the explained variance (Fig. 2C). Besides, PC2 is almost determined by Area and Perimeter with a contribution of 30.9%. Thus, PC1 and PC2 for 2nd screening data also predominantly measure how cells are elongated and enlarged, respectively, in a manner similar to the case of 1st screening. HCA on these PC1 data constructs a dendrogram, showing the presence of two major clusters (Fig. 2D, magenta and orange). As expected, ARHGAP4 and SH3BP1 -which passed 2nd screening -were both located in the identical cluster (magenta).

Alternative HCA was performed based on both the PC1 and PC2 data, showing a similar tendency to the one only on PC1 in that for 2nd screening data all the 15 Rho-GAPs have the same classification as that only on PC1; besides, the alternative one on both the PC1 and PC2 was also found to contain ARHGAP4 and SH3BP1 in the same major cluster (Supplementary Fig. S3). In continuity with our prior study that focused on the extent of cell elongation to detect EMT, which is now found to be predominantly characterized by PC1, we use the PC1 data for the following correlation analyses.

Interestingly, the opposite morphological response was observed between the confluent (1st) and sparse (2nd) conditions. Specifically, cells are elongated compared to control upon the knockdown of the selected 15 Rho-GAPs in confluent culture, while they rather become rounder in sparse culture. The mechanism for this regulation remains elusive, but typically individual epithelial cells in a monolayer (i.e., in confluent culture) display a regular cobble stone-like, relatively round (rather elongated) shape. The silencing of Rho-GAPs would perturb the underlying ordinary balance and result in the loss of the round shape. In contrast, while MCF10A cells in sparse culture typically display an elongated shape, the molecular perturbations to the intact state would result rather in the loss of the elongated shape to turn into a shrank, rounder shape.

### 2.2. Specific members in RhoA-targeting Rho-GAPs share strong correlations between the molecular structural and cell phenotypic similarities

The Rho-GAP domain is an evolutionarily conserved protein domain of the Rho-GAPs, enabling inactivation of the Rho-family with three members of Cdc42, Rac1, and RhoA [18]. Therefore, we further explored if the morphological response of MCF10A cells to the silencing of Rho-GAPs is predictable from the information of the primary structure and the molecular target of the Rho-GAPs [19, 20]. To do this, we quantified the similarity of the cell response (PC1 distance) and that of the amino acid sequence (Structural similarity) between any two of the Rho-GAPs. To determine Structural similarity, Rho-GAP sequences containing all distinguishable domains were collected from UniProt database and analyzed using Clustal Omega. There are 8 different Rho-GAPs, each of which is known to target only RhoA (but not Rac1 and Cdc42) (Fig. 1, 3A). In this case, there are 28 (= _8_C_2_) combinations to select two different Rho-GAPs. For each of the combinations, Structural similarity and PC1 distance were plotted (Supplementary Fig. S4A). Here, two values of Structural similarity were obtained by comparing the “full-length” (Supplementary Fig. S4, red) as well as the “Rho-GAP domain” (Supplementary Fig. S4, blue) of the Rho-GAPs. Likewise, there are 6 Rho-GAPs that target only Rac1, making 15 (= _6_C_2_) combinations (Supplementary Fig. S4B). While there is no Rho-GAP that targets only Cdc42, there are 7 that can be associated with Cdc42 as well as RhoA but not Rac1, making in total 21 (= _7_C_2_) combinations (Supplementary Fig. S4C) that target “RhoA+Cdc42” (but not Rac1). Likewise, there are 13 Rho-GAPs and 78 (= _13_C_2_) combinations that target “Rac1+Cdc42” (but not RhoA) (Supplementary Fig. S4D), and there are 14 Rho-GAPs and 91 (= _14_C_2_) combinations that target “RhoA+Rac1+Cdc42” (Supplementary Fig. S4E). There are 6 other Rho-GAPs whose targets are unclear and thus are not considered in the present analysis.

**Fig. 3.**
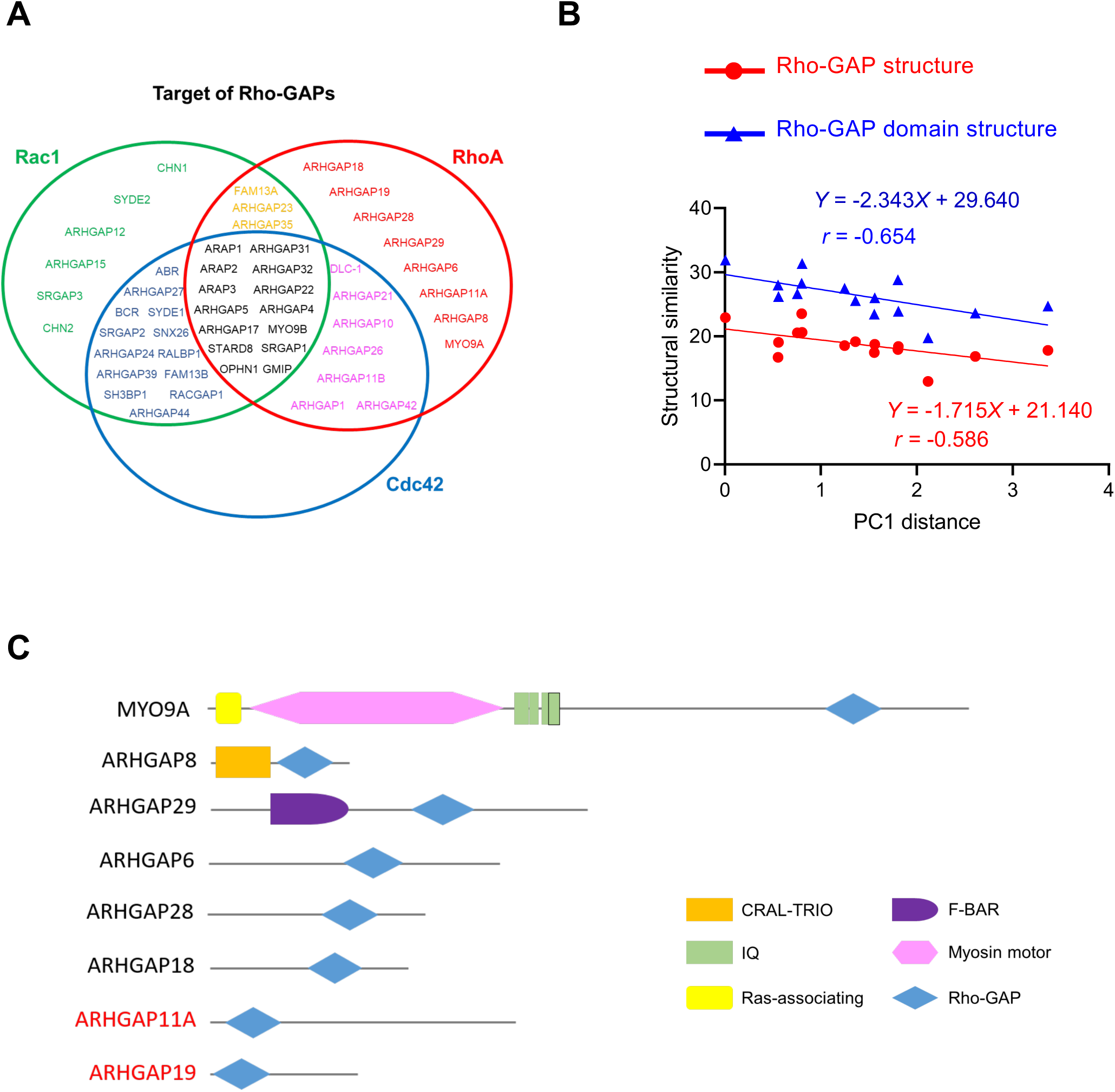
Specific Rho-GAPs share a correlation between the similarity of the primary structure and that of the resulting morphological response of confluent MCF10A cells. (A) Target Rho-family-based categorization of the Rho-GAPs expressed in the cells. 6 Rho-GAPs (specifically, PIK3R1, PIK3R2, INPP5B, DEPDC1, DEPDC1B, OCRL) are not described here because their targets are unclear. (B) The relationship between Structural similarity (considering the full-length structure shown in red or only the Rho-GAP domain structure show in blue) and the PC1 distance -shown in Fig. 2A -of 6 RhoA targeting Rho-GAPs (ARHGAP18, ARHGAP28, ARHGAP29, ARHGAP6, ARHGAP8, and MYO9A). See Supplementary Table S3 for more details. (C) Domain architecture of the 6 Rho-GAPs providing the correlation in B.

For each of the above relationships, linear regression analysis was performed, but we did not see any significant correlation between Structural similarity and PC1 distance, with a small *r* of less than 0.26 in the absolute value (Supplementary Fig. S4). We originally expected that there might be a correlation between them, but given the actual results we alternatively hypothesized that there are some exceptional Rho-GAPs that do not belong to potentially existing correlative relationships. To test this hypothesis, we made a rule that we eliminate at most two molecules from the list of the Rho-GAPs that target only either RhoA (in total 8; Fig. 3A, red font) or Rac1 (in total 6; green font) given the absence of such Rho-GAP affecting only Cdc42; here, importantly, note that all the possible combinations of the one or two molecules to be eliminated were tested, meaning that this procedure is conducted totally in an objective way. We thus examined all the combinations and consequently found that, if we remove two Rho-GAPs that target only RhoA -specifically, ARHGAP11A and ARHGAP19 -, it turns out that the rest of the 6 (ARHGAP18, ARHGAP28, ARHGAP29, ARHGAP6, ARHGAP8, and MYO9A; i.e., 15 (= _6_C_2_) combinations, Fig. 3C) possess a level of correlation, between Structural similarity and PC1 distance, of an *r* of -0.586 for the full-length of the Rho-GAPs (Fig. 3B, red) or -0.654 for the Rho-GAP domain of the Rho-GAPs (Fig. 3B, blue; Supplementary Table S3). These negative values -strong in correlation magnitude according to Evans’ classification [21] -indicate that the more similar their primary structures are, the more similar change cells undergo following the silencing of the Rho-GAPs.

On the other hand, any removal of the two Rho-GAPs that target only Rac1 did not improve the correlation significantly. At best, removal of CHN1 and CHN2 results in a slight improvement in *r* up to -0.126 for the full-length group and -0.158 for the Rho-GAP domain group (figure not shown). In addition, removal of only one Rho-GAP that targets only RhoA or Rac1 was not enough for allowing the rest to have a correlation (data not shown). Together, we detected the presence of a strong correlation between the protein structural and cell morphological similarities only in the specific members of RhoA-targeting Rho-GAPs (but not Rac1-targeting ones) excluding ARHGAP11A and ARHGAP19. We will discuss below possible reasons for why the removal of the two Rho-GAPs led to the emergence of the strong correlation; again, note that we here uncovered this specific combination based on the completely objective way since the removal of any one was not enough, whereas the removal of two (tested for any combination) reached the strong correlation.

We also focused on the 15 Rho-GAPs (Fig. 2D) that passed 1st screening to see if there are such correlations for sparsely plated cells as well. The specific target was unclear regarding PIK3R1, DEPDC1B, and OCRL, and therefore the remaining 12 were analyzed. Because they do not have the only one target except for ARHGAP11A that targets only RhoA, we considered 3 groups in which the constituents have a target with at least one of RhoA, Rac1, or Cdc42 (Supplementary Fig. S5A). Linear regression analyses for all the 3 groups, however, show no significant correlation between Structural similarity and PC1 distance with at most an *r* of -0.17 for Rho-GAPs that can target RhoA (Supplementary Fig. S5A; Supplementary Table S4). We also separately confirmed that RhoA-targeting Rho-GAPs enclose, on average, a smaller area than Rac1- and Cdc42-related ones do in the PC1-PC2 coordinate plane (Supplementary Fig. S5B), being in line with the above conclusion that RhoA-targeting Rho-GAPs tend to exhibit a similar response in the cell morphology.

### 2.3. Change in cell shape due to the silencing of specific Rho-GAPs is correlated with that in migratory ability

We next investigated the relationship, which sparsely plated cells may have, between the migration ability and morphological response to the silencing of Rho-GAPs. We focused on a major cluster that contains ARHGAP4 and SH3BP1 that are particularly responsible for maintaining the cell morphology in both confluent and sparse cultures [13]. Migration speed of single cells subjected to the silencing of one of the 7 Rho-GAPs (Fig. 2D, magenta) was evaluated (Supplementary Fig. S6), in which two different siRNAs were used for each case including control. We found that the silencing of each of ARHGAP4, ARHGAP11A, ARHGAP17, and SH3BP1 all decreases the migration speed significantly (Fig. 4A). One type of siRNA for MYO9B knockdown decreased the migration speed significantly as reported previously [22], but in the present case the tendency was not consistent over different siRNAs.

**Fig. 4.**
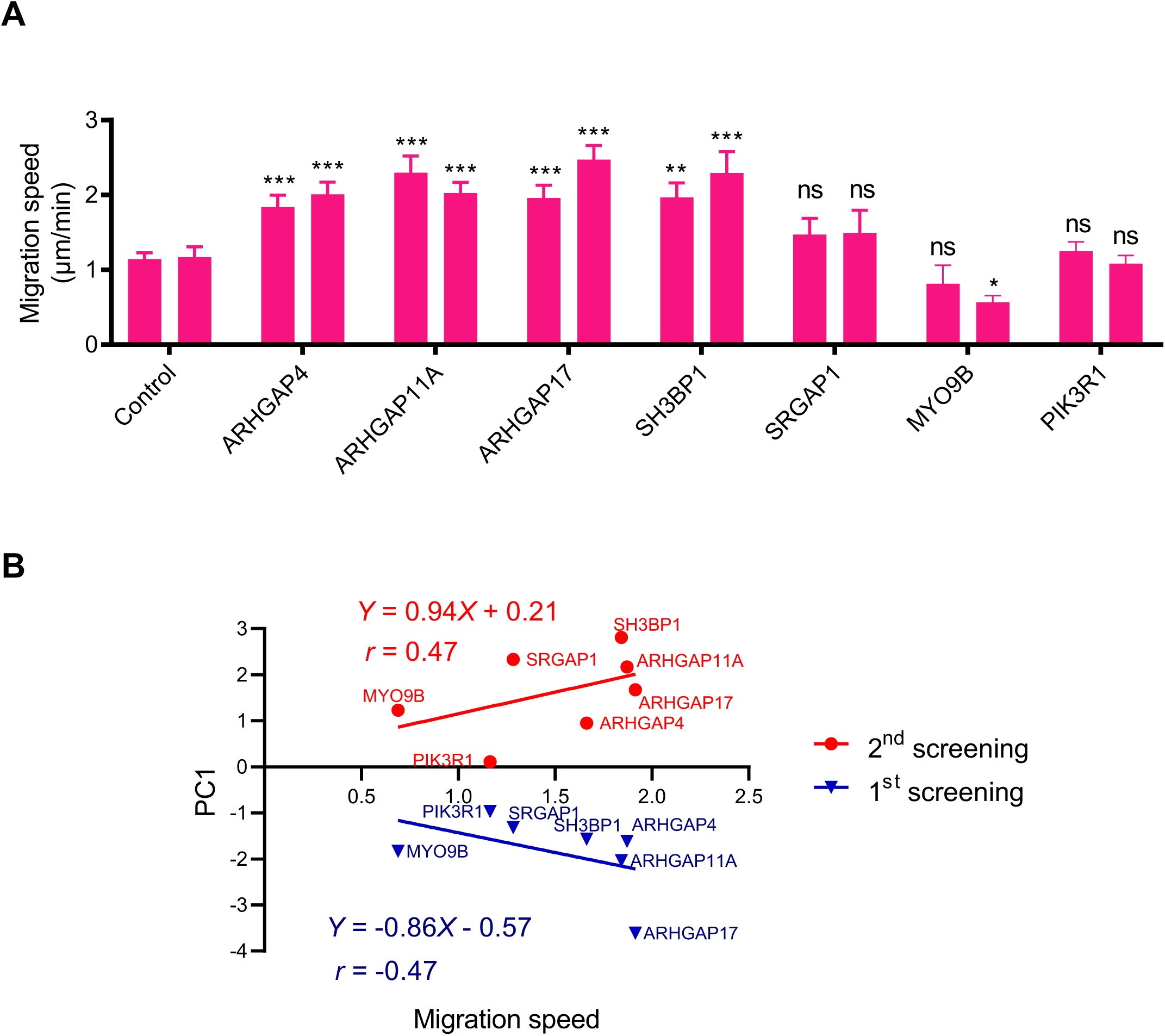
Rho-GAPs that induce significant morphological response in MCF10A cells also tend to increase the migratory potential. (A) Migration speed examined for sparsely plated individual cells that are subject to the silencing of the Rho-GAPs categorized in HCA in the identical cluster (Fig. 2D, magenta). Two siRNAs were tested, and for each case the data were obtained from 3 independent experiments with *n* ≥ 10 cells. (B) PC1 value for 1st (blue) and 2nd (red) screening data as a function of Migration speed of cells subjected to the silencing of the specific Rho-GAP indicated.

The mean of the migration speed was normalized for each case by the mean of the controls and was plotted as a function of their respective PC1 values obtained in sparse culture (Fig. 2C), suggesting the presence of a moderate correlation with an *r* of 0.47 (Fig. 4B, red). The positive *r* value indicates that the extent of how much cells become rounder in shape upon the silencing of the clustered 7 Rho-GAPs is positively correlated with that of how much cells are enhanced in migratory ability. If we instead use PC1 values obtained in confluent culture (Fig. 2A), a correlation similar in magnitude but opposite in sign of an *r* of -0.47 was detected (Fig. 4B, blue), suggesting that the extent of elongation of cells in monolayers upon the knockdown is negatively correlated with the simultaneous enhancement of the migratory ability (Fig. 4B, blue).

## 3. Discussion

Rho-family -with three members of Cdc42, Rac1, and RhoA -is regulated in a highly complicated manner by a large number of upstream pathways to mediate a wide variety of cell functions [1-3]. Among them, in humans there are 66 Rho-GAPs that potentially control Rho-family. While the involvement of specific Rho-GAPs in cell functions (so far mostly regarding cell migration) has gradually been elucidated, comprehensive understanding of that of the whole family is currently quite far from complete. Motivated by this situation, here we investigated to see if there is a consistent relationship between specific Rho-GAPs and cell phenotypes; in other words, a certain consequent phenotype -specifically on the morphology and migratory potential -might be predictable from the molecular structure of Rho-GAPs. We found that, while absent over the entire family, if we focus on specific RhoA-targeting Rho-GAPs there is a strong correlation between the cell morphology and primary structure of the molecule. This finding is in line with that in previous studies reporting the importance of RhoA in the regulation of cell morphology [23].

Of all the 57 Rho-GAPs expressed in human epithelial MCF10A cells, 8 Rho-GAPs are known to inactivate the GTPase activity of RhoA but not that of Rac1 nor Cdc42. We did not find significant correlations regarding all the 8, but if we eliminate two -ARHGAP11A and ARHGAP19 -from them, then the rest exhibit a strong correlation between the similarities of the molecular structure and of epithelial cell response to the molecular silencing for any two from the remaining 6 Rho-GAPs. This finding prompts us to conclude that the target and structure of Rho-GAPs are critical factors to determine the manner of morphological alterations in MCF10A cells forming a confluent monolayer. As the correlation is greater in magnitude for the case limiting only to the Rho-GAP domain of the Rho-GAPs for the quantification of Structural similarity (Fig. 3B, blue) compared to the one considering the full-length structure (red), the sequence of the Rho-GAP domain might be a major determinant for the expressed cellular response [24].

The above 6 RhoA-targeting Rho-GAPs include ARHGAP18, ARHGAP28, ARHGAP29, ARHGAP6, ARHGAP8, and MYO9A (Fig. 3C). Among them, the silencing of ARHGAP18 has been reported to induce a rounder shape in HeLa cancer cells [25], i.e., consistent with our observations on epithelial MCF10A cells plated in sparse culture (Fig. 2C). There is a report, though without quantification, that the silencing of MYO9A in human epithelial intestinal Caco-2 cells alters the cell morphology and junctions [26], while we observed only a moderate change in MCF10A cells (Fig. 2A). On the other hand, we could not find literature describing the role of the rest (ARHGAP28, ARHGAP29, ARHGAP6, or ARHGAP8) in the regulation of cell morphology. To allow the relationship to have the strong correlation, we had to eliminate ARHGAP11A and ARHGAP19. Unlike the above-described 6, these two are the closest to each other in terms of Structural similarity, containing a Rho-GAP domain located close to the N-terminus with no other distinguishable domain in between the N-terminus and the Rho-GAP domain (Fig. 3C). These structures might have some effect -which the other 6 do not - to be excluded from the group that shares the correlative relationship. One possible explanation might be that ARHGAP11A, which targets only RhoA, has been reported to directly interact with Rac1B in a manner independent of the Rho-GTPase activity [27]. As a result, ARHGAP11A is involved in both EMT and its reverse mesenchymal-epithelial transition (MET) by increasing the activities of Rac1B and RhoA, respectively. Given this different mechanism of the interaction with Rho-family, the exclusion of ARHGAP11A from the correlating group, which is assured by the GTPase activity-dependent uniqueness of the target, might make sense. Meanwhile, ARHGAP19 has been reported to target miR-200c that is known to be involved in EMT and hence cell morphological changes [28]. Thus, ARHGAP19 might also more or less affect the cell morphology predominantly in a mechanism distinct from the correlating group using the Rho-GTPase activity.

By analyzing the primary cluster in 2nd screening (Fig. 2C, magenta), we also demonstrated that 4 Rho-GAPs -ARHGAP4, ARHGAP11A, ARHGAP17, and SH3BP1 -all affect the migration of sparsely plated MCF10A cells, while the rest 3 -SRGAP1, MYO9B, and PIK3R1 -do not (Fig. 4A). These 7 were detected in 1st screening as a factor that significantly elongates the morphology of confluent cells upon the silencing (Fig. 2A). Meanwhile, their effects on the morphology of sparse cells were diverse in magnitude, but on average the silencing gave rise to a rounder shape (Fig. 2C). There is thus a variation in the cell response, but interestingly we found another distinct tendency that the 7 Rho-GAPs create in cells between the morphological response (PC1 value) and migration speed under the silencing of the Rho-GAPs. This tendency persists regardless of the source (1st or 2nd screening data) of PC1 values and hence may provide a solid foundation for comprehensive understanding of the role of Rho-GAPs. Particularly, the presence of the correlation may suggest a general feature that morphological change and increased migration of epithelial cells are complementary processes to mediate EMT.

## 4. Materials and methods

### 4.1. Rho-GAP screening on cell morphology

We previously analyzed the effect of silencing Rho-GAPs in human epithelial cells [13]. Briefly, while at least 66 genes are known to encode Rho-GAPs, we identified that 57 Rho-GAPs are expressed in MCF10A human mammary epithelial cells (ATCC, Manassas, USA). For each of these, the effect of siRNA-based knockdown on the morphology of individual cells was evaluated by manually extracting, with the greatest care, the geometric information of the outline of the cells in ImageJ software (NIH, Bethesda, USA) using an electric pen device (Cintiq 13HD, DTK-1301, Wacom, Japan) from two independent experiments, in each of which at least 1,000 cells were analyzed for each group [13] (Supplementary Fig. S7). We then identified 15 Rho-GAPs that play a significant role in maintaining a round cell shape (Fig. 1, red; 1st screening). We performed the same experiments on the extracted 15 Rho-GAPs in sparse cells to finally identify 2 Rho-GAPs as a major regulator maintaining a round cell shape (Fig. 1, blue; 2nd screening).

### 4.2. Principal components analysis and hierarchical cluster analysis

In the above screening, the decreased roundness of individual cell shape -i.e., a measure of EMT -was evaluated using ImageJ based on two morphological parameters: Circularity and Aspect ratio (AR). Circularity is defined as 4π × (Area)/(Perimeter)^2^, being 1 for a circular shape and 0 for a linear one. AR represents the ratio of the long axis to the vertical axis of cells approximated as an elliptical shape. To more thoroughly evaluate the effect of the knockdown on the cell morphology in the present study, three more parameters were additionally considered for the same dataset: Area, Perimeter, and Solidity. The former two were already included in the calculation of Circularity, but here they are explicitly and independently evaluated. Solidity is another parameter calculated in ImageJ once the outline of the cell is determined, which represents the area divided by the convex area (an imaginary convex hull around it); consequently, it gets close to 1 with a smooth morphology while gets smaller with a tortuous or branched morphology. Regarding these 5 parameters (normalized by dividing the mean by the mean of siControl conditions), principal components analysis (PCA) was performed using Origin (OriginLab Corporation, Northampton, USA) to obtain PC1 and PC2, i.e., the specific directions in the 5-dimensional data space along which the data display the largest and second largest variations, respectively [29]. Hierarchical cluster analysis (HCA) was then performed using Ward linkage method based on the PC1 data or both PC1 and PC2 data [30]. PCA and HCA were performed for both cases of 1st (57 Rho-GAPs) and 2nd (15 Rho-GAPs) screening.

### 4.3. Target gene contribution analysis

In the above result of PCA/HCA for 1st screening, Rho-GAPs can be divided into several major clusters. The area of a planar polygon within the PC1-PC2 coordinate system enclosed by the Rho-GAPs contained in the identical clusters and having the same molecular target in terms of the GTPase activity (i.e., RhoA, Cdc42, and Rac1) is then calculated. Here, to determine the target(s) of each Rho-GAP in terms of the GTPase activity, we referred to a previous report [31]. It is expected that the smaller the enclosed area is, the more similar is the effect of the Rho-GAPs on the morphological response of cells. We also investigated if there is a specific tendency in the relationship between the primary structure and PC1 among different Rho-GAPs. The amino acid sequence of the “full-length” or “Rho-GAP domain” of Rho-GAPs found in the UniProt database was submitted to Clustal Omega (https://www.ebi.ac.uk/Tools/msa/clustalo/) to obtain a matching score between any two of them [32, 33]. The matching score, which ranges between 0% (for the lowest similarity) and 100% (for the highest similarity), is here called Structural similarity and plotted as a function of the Euclidean distance between PC1 values of two Rho-GAPs, which we term PC1 distance. The relationship between Structural similarity and PC1 distance is examined considering the molecular target(s) of the Rho-GAPs.

### 4.4. Cell migration assay

A cell suspension was added to each well of a 24-well plate and incubated for 24 hours. Single cells were chosen and imaged every 10 minutes for up to 2 days using a camera (ORCA-R2, Hamamatsu Photonics, Hamamatsu, Japan) on a bright-field microscope (IX73, Olympus, Tokyo, Japan). The central position of individual cells was automatically determined by ImageJ once the outline of the cells was identified (Supplementary Fig. S6). The migration velocity of cells was then determined by analyzing the total length (but not the end-to-end distance between the initial and final time points) that the cells migrate over the total time period. More than 10 cells from more than 3 independent experiments were assessed for each Rho-GAP group.

### 4.5. Statistical analysis

Statistical analysis was performed using Prism 8 (GraphPad Software, San Diego, USA), in which linear regression lines and Pearson correlation coefficients *r* were obtained by the least-squares method. Data for the cell migration were shown as mean ± standard error of mean from more than three independent experiments. Statistical significance compared to siControl was set as follows: *, p < 0.05; ***, p < 0.001.

## Acknowledgment

K.N. is supported by Chinese Scholarship Council (CSC) Scholarship.

## Dada availability statement

The data that support the findings of this study are available from the corresponding author upon reasonable request.

## Author contributions

K.N., T.S.M., and S.D. designed the research; K.N. performed the experiments and analyses; T.S.M. and S.D. provided technical support; K.N. and S.D. wrote the paper. All the authors provided editorial comments and approved the manuscript.

## Supplementary materials

### List of the Supplementary materials included

#### Supplementary Figures

**Fig. S1.**
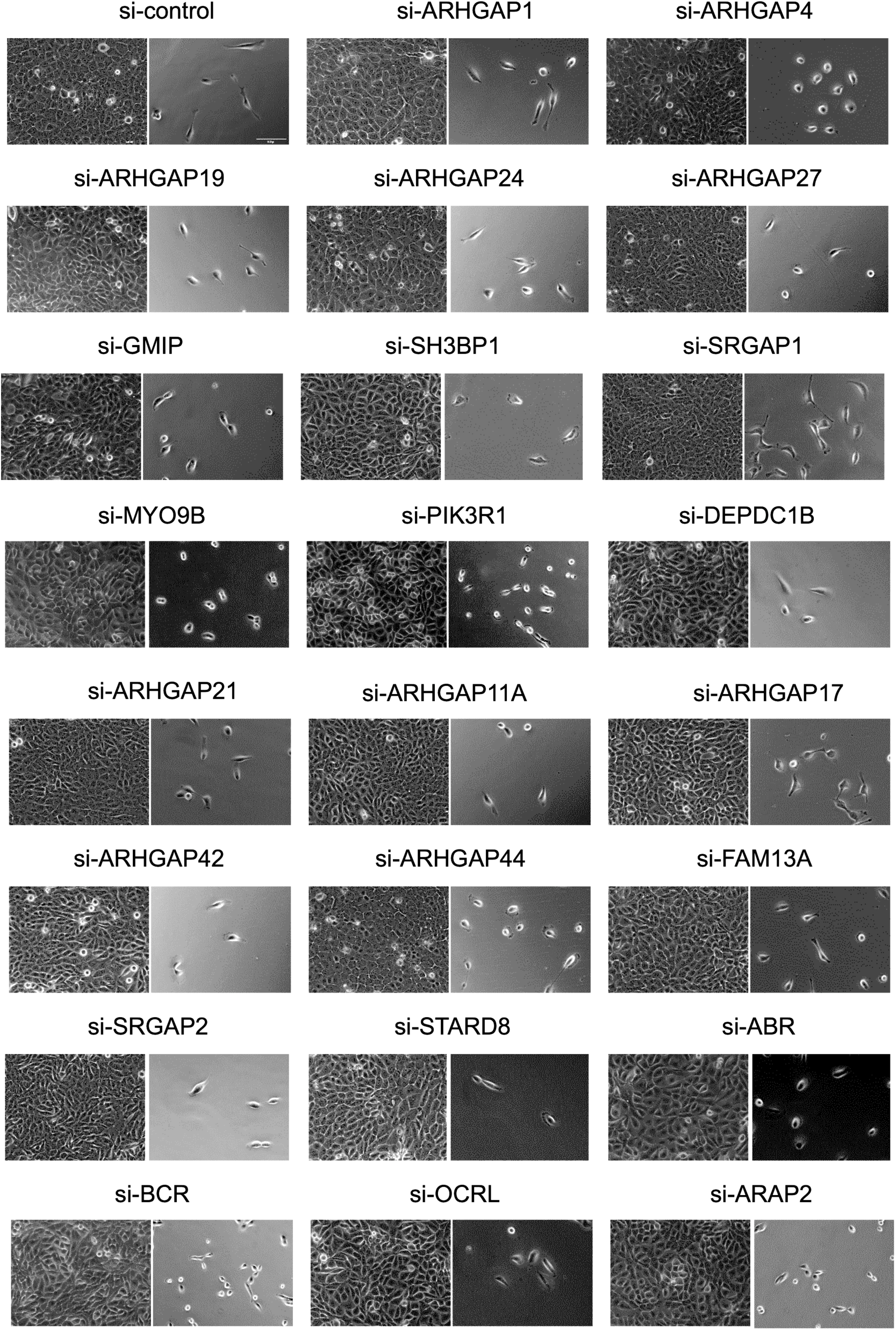
Phase-contrast images of MCF10A cells in confluent (left) and sparse (right) culture subjected to the siRNA-induced silencing of the Rho-GAP indicated. Scale, 100 μm. For this cell screening experiment, MCF10A human mammary epithelial cells (ATCC, Manassas, USA) were cultured in DMEM/Ham’s F12 medium (Wako, Osaka, Japan) supplemented with 5% horse serum (Gibco, Waltham, USA), 0.25 IU/ml insulin (Wako), 0.5 mg/ml hydrocortisone (Wako), 100 ng/ml cholera toxin (Wako), 20 ng/ ml EGF (Wako), and 1% penicillin-streptomycin solution (Wako) in a 5% CO_2_ incubator at 37°C. Cells were transfected with each of the siRNAs targeting the human Rho-GAPs or negative control siRNAs (Thermo Fisher Scientific, Waltham, USA) using Lipofectamine RNAiMAX Reagent (Thermo Fisher Scientific) according to the manufacturer’s instructions. Cells were imaged, 36 h after the transfection, using an ORCA-R2 camera (Hamamatsu Photonics, Hamamatsu, Japan) on a phase-contrast microscope (IX73, Olympus, Tokyo, Japan).

**Fig. S2.**
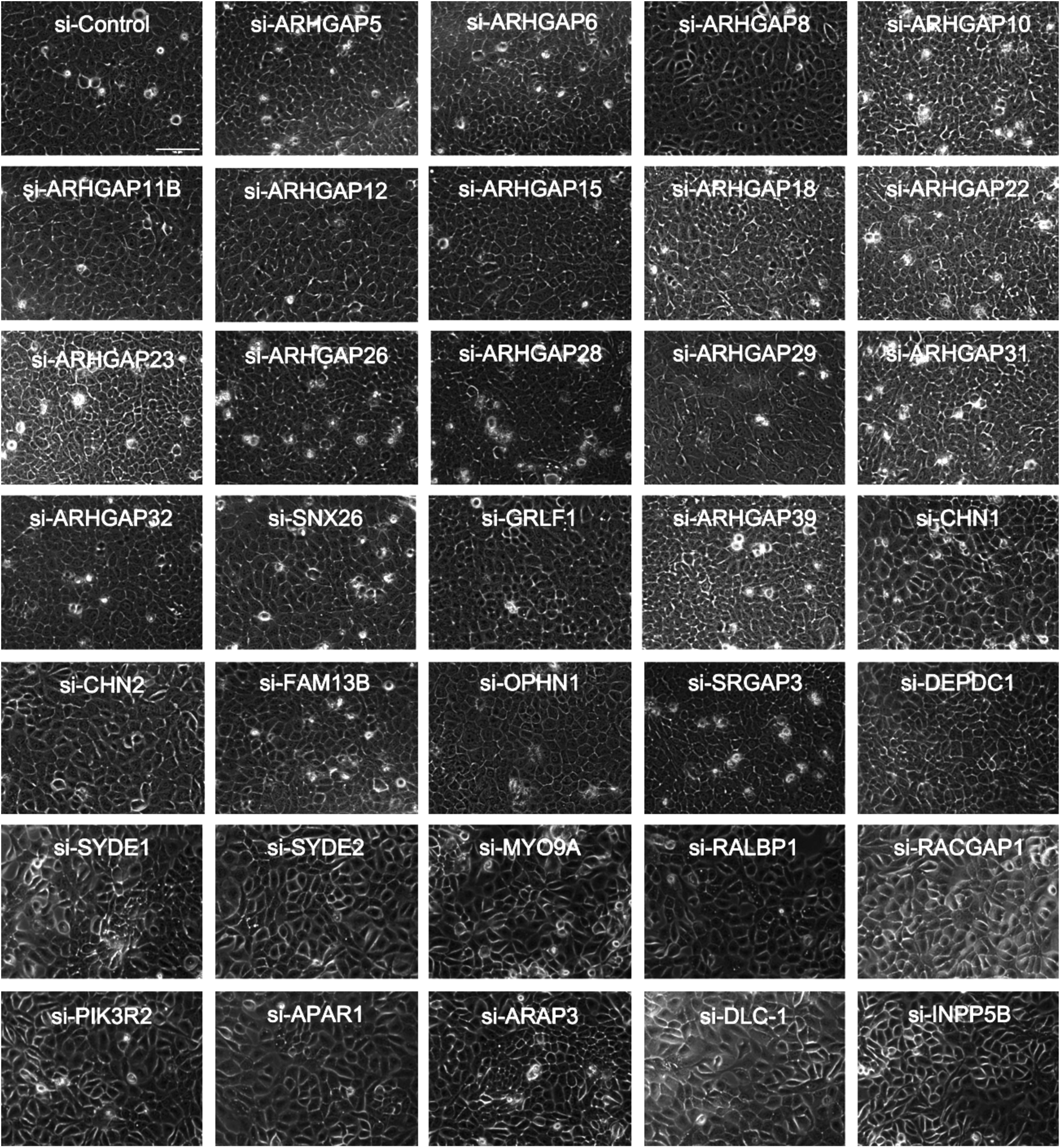
Phase-contrast images of MCF10A cells in confluent culture subjected to the siRNA-induced silencing of the Rho-GAP indicated. Scale, 100 μm.

**Fig. S3.**
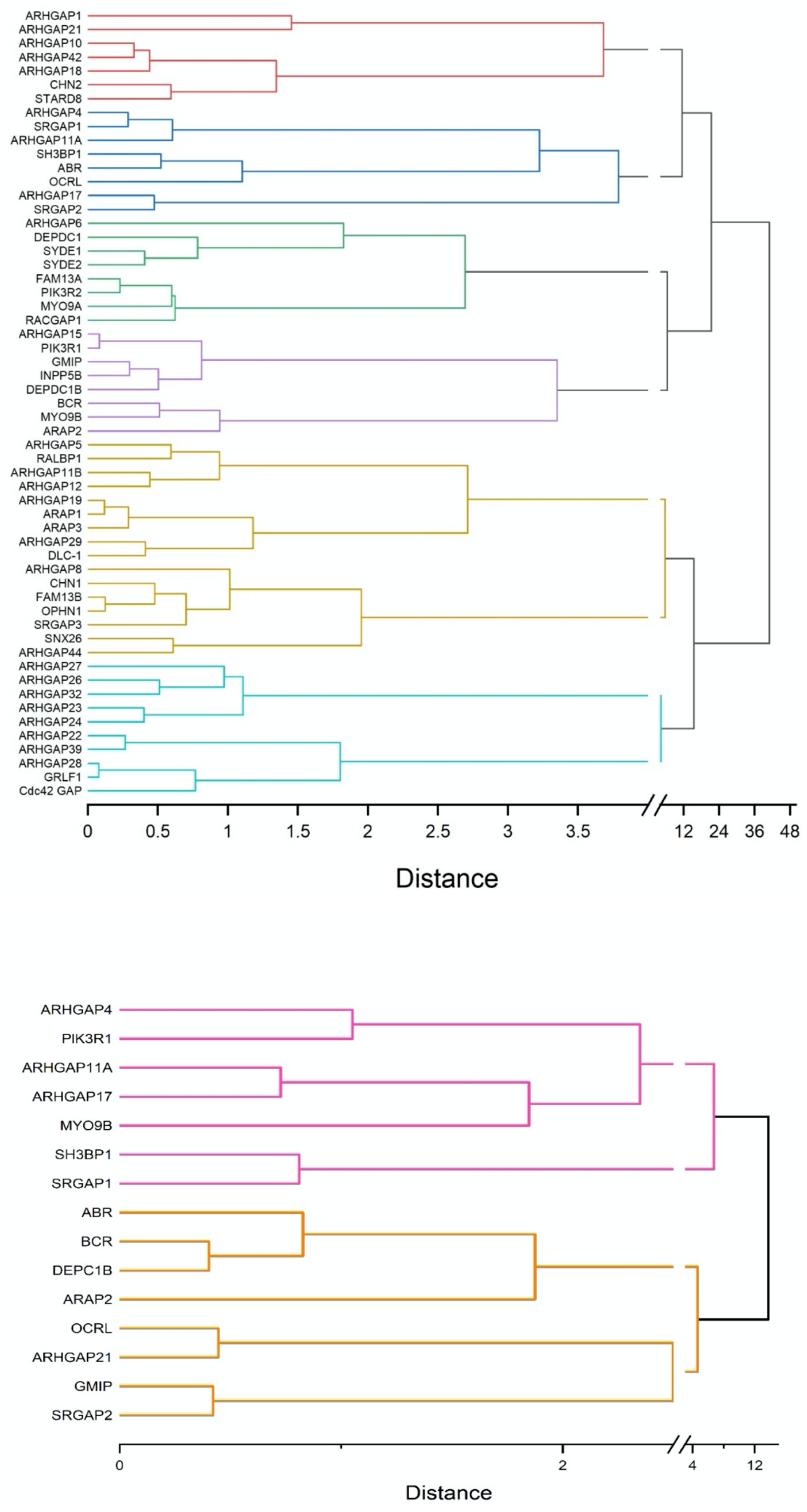
HCA based on both PC1 and PC2 for 1st (upper) and 2nd (lower) screening data.

**Fig. S4.**
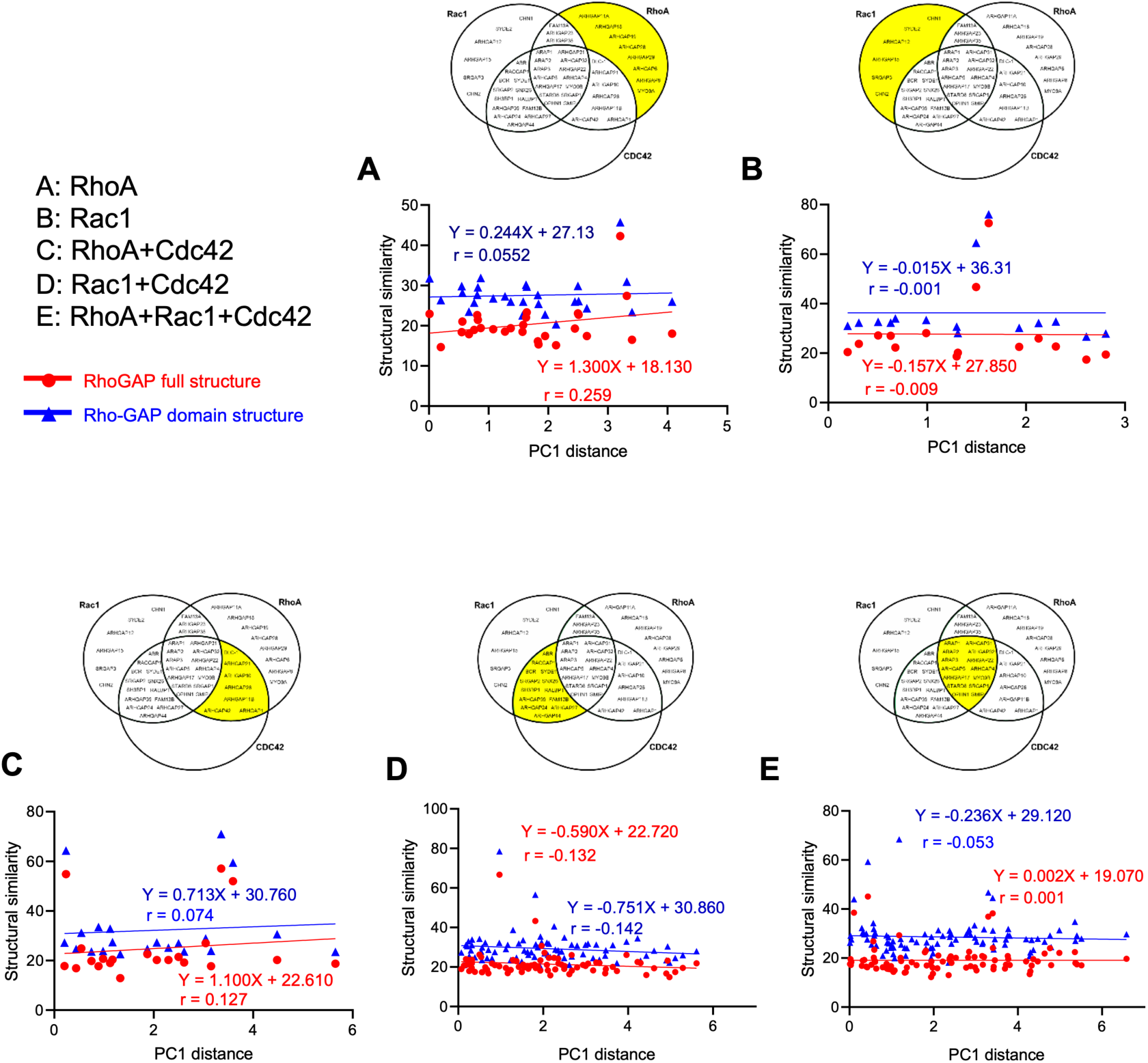
The relationship between Structural similarity and PC1 distance for confluent cells. All the pair selected from the Rho-GAPs in the same target group of A–E were analyzed. The group examined is highlighted in yellow in the Venn diagrams.

**Fig. S5.**
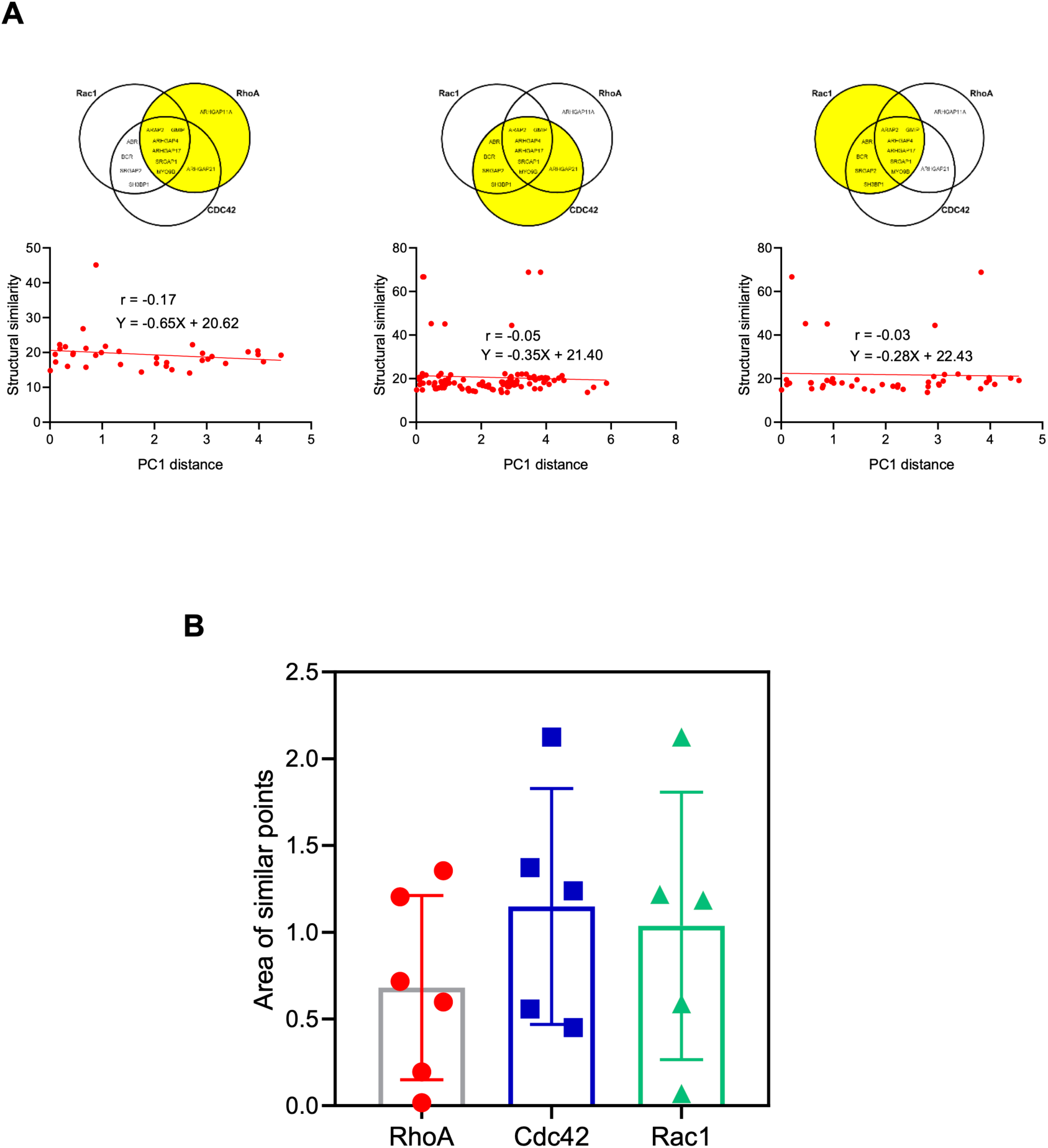
Additional analyses to find a possible specific relationship among the Rho-GAP structure, target, and the cell morphological response. (A) The relationship between Structural similarity and PC1 distance for sparse cells. All the pair selected from the Rho-GAPs in the same target group (highlighted in yellow in the Venn diagrams) were analyzed. See Table S3 for more details. (B) The area of a planar polygon within the PC1-PC2 coordinate system for confluent cells (Fig. 2A) enclosed by the specific Rho-GAPs contained in the identical clusters (Fig. 2B) is grouped depending on the GTPase activity target (RhoA, Cdc42, or Rac1). Data are shown with plots and the mean ± standard deviation.

**Fig. S6.**
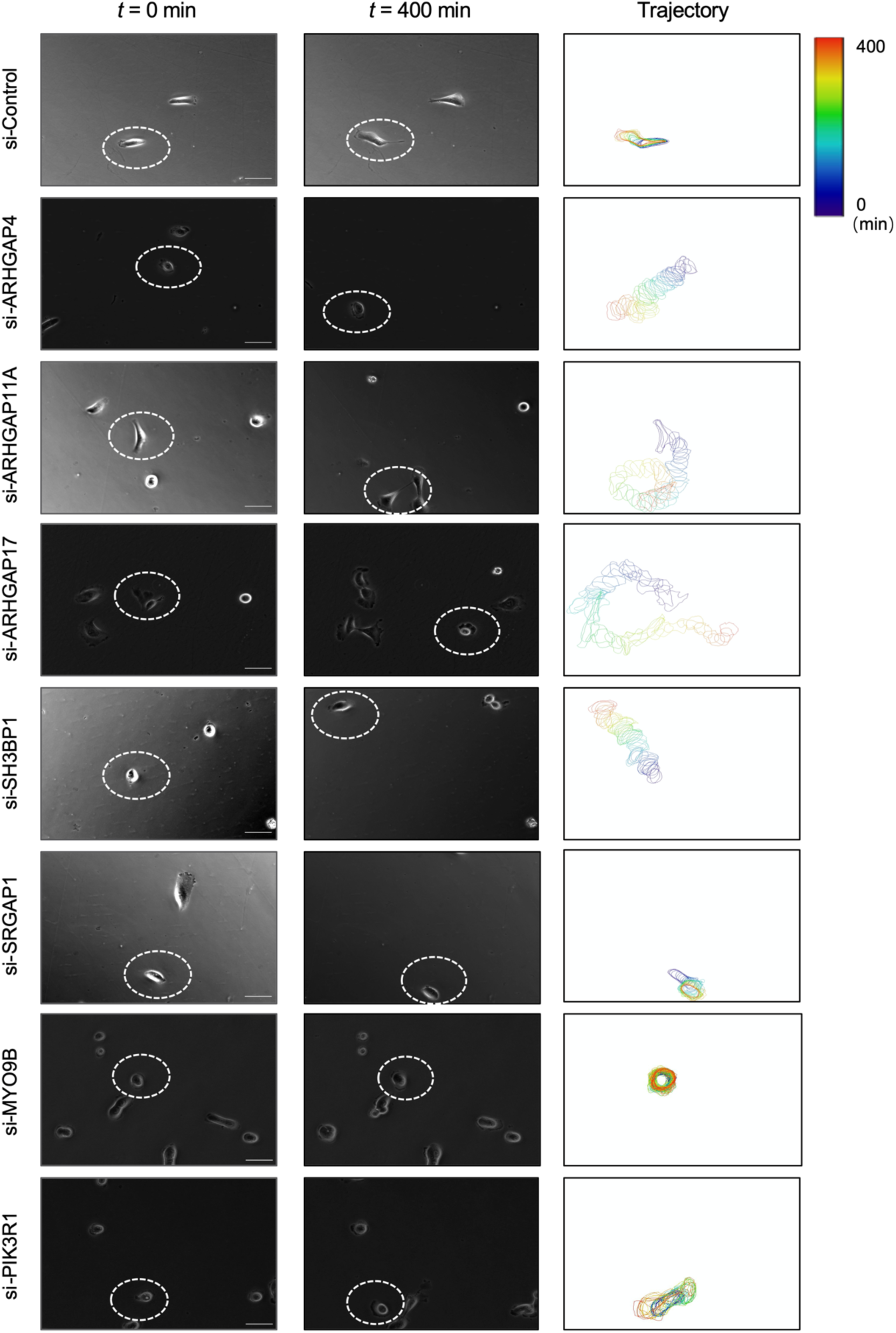
Trajectory extraction of cell migration subjected to the silencing of siControl or one of the 7 Rho-GAP siRNAs. Scale, 50 μm.

**Fig. S7.**
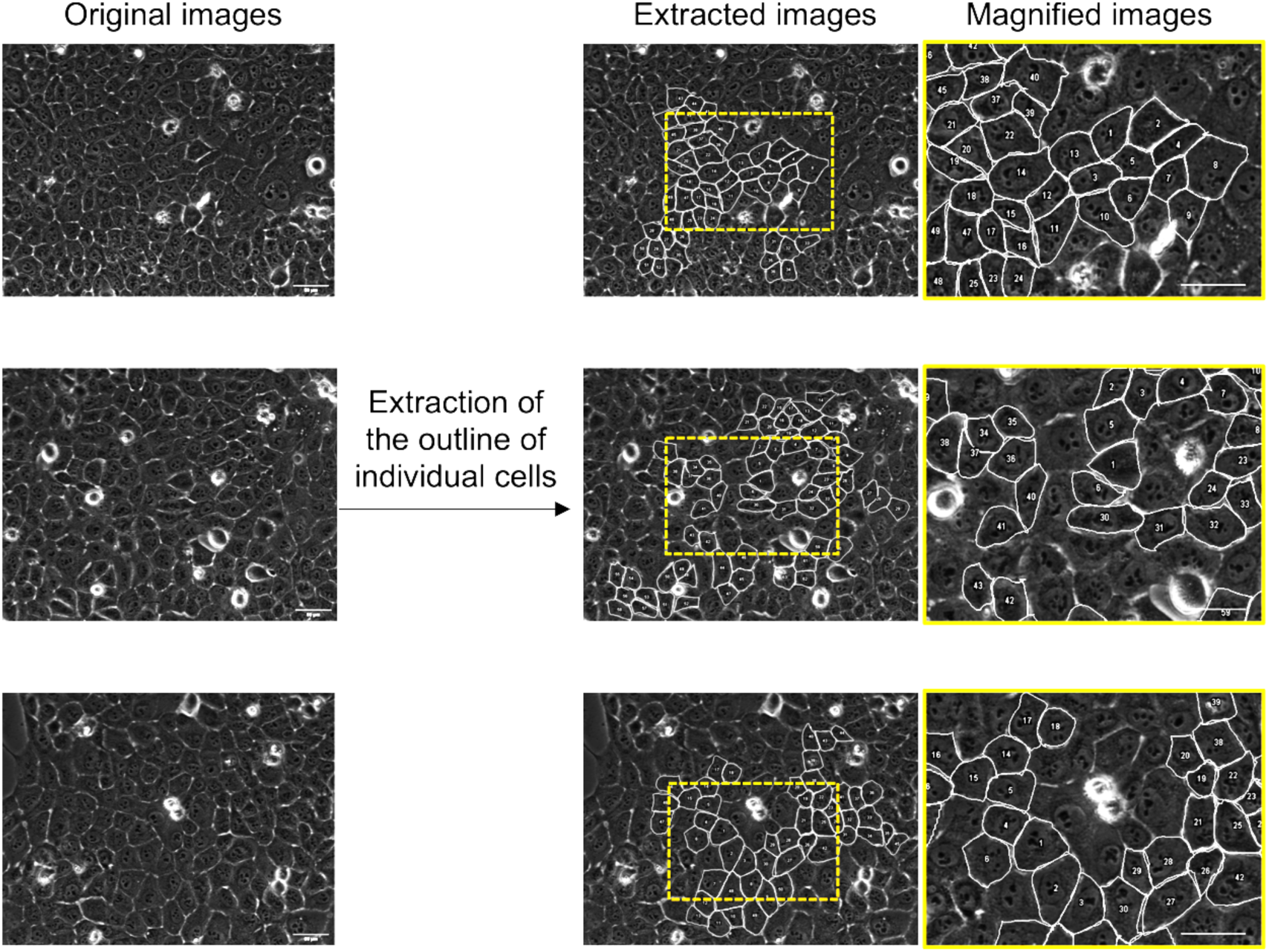
Examples of the extraction of cell morphology (siControl) from two independent experiments, in each of which the data are comprised of at least 1,000 cells for each group (see “Materials and methods” and “Dada availability statement” for details). Scale, 50 μm.

#### Supplementary Tables

Table S1 Numerical data of Fig. 2A, with the Rho-GTPase activity target(s) of the Rho-GAPs.

Table S2 Numerical data of Fig. 2C.

Table S3 Structural similarity between the two Rho-GAPs analyzed in Fig. 3B.

Table S4 Structural similarity between the two Rho-GAPs analyzed in Fig. S5A.

## References

1. Fujiwara, S., Matsui, T. S., Ohashi, K., Deguchi, S., Mizuno, K. 2018 Solo, a RhoA-targeting guanine nucleotide exchange factor, is critical for hemidesmosome formation and acinar development in epithelial cells. PLoS One. 13, e0195124. (10.1371/journal.pone.0195124)

2. Etienne-Manneville, S., Hall, A. 2002 Rho GTPases in cell biology. Nature. 420, 629-635. (10.1038/nature01148)

3. Haga, R. B., Ridley, A. J. 2016 Rho GTPases: Regulation and roles in cancer cell biology. Small GTPases. 7, 207-221. (10.1080/21541248.2016.1232583)

4. Ahmed, S., Bu, W., Lee, R. T. C., Maurer-Stroh, S., Goh, W. I. 2010 F-BAR domain proteins: Families and function. Commun Integr Biol. 3, 116-121. (10.4161/cib.3.2.10808)

5. Panagabko, C., Morley, S., Hernandez, M., Cassolato, P., Gordon, H., Parsons, R., Manor, D., Atkinson, J. 2003 Ligand specificity in the CRAL-TRIO protein family. Biochemistry. 42, 6467-6474. (10.1021/bi034086v)

6. Zinatizadeh, M. R., Momeni, S. A., Zarandi, P. K., Chalbatani, G. M., Dana, H., Mirzaei, H. R., Akbari, M. E., Miri, S. R. 2019 The Role and Function of Ras-association domain family in Cancer: A Review. Genes Dis. 6, 378-384. (10.1016/j.gendis.2019.07.008)

7. Vega, F. M., Ridley, A. J. 2008 Rho GTPases in cancer cell biology. FEBS Lett. 582, 2093-2101. (10.1016/j.febslet.2008.04.039)

8. Zuo, Y., Oh, W., Ulu, A., Frost, J. A. 2016 Minireview: Mouse Models of Rho GTPase Function in Mammary Gland Development, Tumorigenesis, and Metastasis. Molecular Endocrinology. 30, 278-289. (10.1210/me.2015-1294)

9. van Buul, J. D., Geerts, D., Huveneers, S. 2014 Rho GAPs and GEFs. Cell Adhesion & Migration. 8, 108-124. (10.4161/cam.27599)

10. Xu, J., Zhou, X. L., Wang, J. L., Li, Z. L., Kong, X., Qian, J., Hu, Y., Fang, J. Y. 2013 RhoGAPs Attenuate Cell Proliferation by Direct Interaction with p53 Tetramerization Domain. Cell Reports. 3, 1526-1538. (10.1016/j.celrep.2013.04.017)

11. Amin, E., Jaiswal, M., Derewenda, U., Reis, K., Nouri, K., Koessmeier, K. T., Aspenstrom, P., Somlyo, A. V., Dvorsky, R., Ahmadian, M. R. 2016 Deciphering the Molecular and Functional Basis of RHOGAP Family Proteins: A SYSTEMATIC APPROACH TOWARD SELECTIVE INACTIVATION OF RHO FAMILY PROTEINS. Journal of Biological Chemistry. 291, 20353-20371. (10.1074/jbc.M116.736967)

12. Muller, P. M., Rademacher, J., Bagshaw, R. D., Wortmann, C., Barth, C., van Unen, J., Alp, K. M., Giudice, G., Eccles, R. L., Heinrich, L. E., et al. 2020 Systems analysis of RhoGEF and RhoGAP regulatory proteins reveals spatially organized RAC1 signalling from integrin adhesions. Nat Cell Biol. 22, 498-511. (10.1038/s41556-020-0488-x)

13. Kang, N., Matsui, T. S., Liu, S., Fujiwara, S., Deguchi, S. 2020 Comprehensive analysis on the whole Rho-GAP family reveals that ARHGAP4 suppresses EMT in epithelial cells under negative regulation by Septin9. FASEB J. 10.1096/fj.201902750RR)

14. Blangy, A. 2017 Tensins are versatile regulators of Rho GTPase signalling and cell adhesion. Biol Cell. 109, 115-126. (10.1111/boc.201600053)

15. Lawson, C. D., Fan, C., Mitin, N., Baker, N. M., George, S. D., Graham, D. M., Perou, C. M., Burridge, K., Der, C. J., Rossman, K. L. 2016 Rho GTPase Transcriptome Analysis Reveals Oncogenic Roles for Rho GTPase-Activating Proteins in Basal-like Breast Cancers. Cancer Research. 76, 3826-3837. (10.1158/0008-5472.Can-15-2923)

16. Leggett, S. E., Sim, J. Y., Rubins, J. E., Neronha, Z. J., Williams, E. K., Wong, I. Y. 2016 Morphological single cell profiling of the epithelial-mesenchymal transition. Integr Biol (Camb). 8, 1133-1144. (10.1039/c6ib00139d)

17. Shen, L., Qu, X., Ma, Y., Zheng, J., Chu, D., Liu, B., Li, X., Wang, M., Xu, C., Liu, N., et al. 2014 Tumor suppressor NDRG2 tips the balance of oncogenic TGF-β via EMT inhibition in colorectal cancer. Oncogenesis. 3, e86-e86. (10.1038/oncsis.2013.48)

18. Barrett, T., Xiao, B., Dodson, E. J., Dodson, G., Ludbrook, S. B., Nurmahomed, K., Gamblin, S. J., Musacchio, A., Smerdon, S. J., Eccleston, J. F. 1997 The structure of the GTPase-activating domain from p50rhoGAP. Nature. 385, 458-461. (10.1038/385458a0)

19. Peck, J., Douglas, G. t., Wu, C. H., Burbelo, P. D. 2002 Human RhoGAP domain-containing proteins: structure, function and evolutionary relationships. FEBS Lett. 528, 27-34. (10.1016/s0014-5793(02)03331-8)

20. Tcherkezian, J., Lamarche-Vane, N. 2007 Current knowledge of the large RhoGAP family of proteins. Biol Cell. 99, 67-86. (10.1042/BC20060086)

21. Evans, J. D. 1996 Straightforward Statistics for the Behavioral Sciences. Brooks/Cole Publishing Company.

22. Hanley, P. J., Xu, Y., Kronlage, M., Grobe, K., Schon, P., Song, J., Sorokin, L., Schwab, A., Bahler, M. 2010 Motorized RhoGAP myosin IXb (Myo9b) controls cell shape and motility. Proc Natl Acad Sci U S A. 107, 12145-12150. (10.1073/pnas.0911986107)

23. Kitzing, T. M., Sahadevan, A. S., Brandt, D. T., Knieling, H., Hannemann, S., Fackler, O. T., GroBhans, J., Grosse, R. 2007 Positive feedback between Dia1, LARG, and RhoA regulates cell morphology and invasion. Gene Dev. 21, 1478-1483. (10.1101/gad.424807)

24. Amin, E., Jaiswal, M., Derewenda, U., Reis, K., Nouri, K., Koessmeier, K. T., Aspenström, P., Somlyo, A. V., Dvorsky, R., Ahmadian, M. R. 2016 Deciphering the Molecular and Functional Basis of RHOGAP Family Proteins. Journal of Biological Chemistry. 291, 20353-20371. (10.1074/jbc.M116.736967)

25. Maeda, M., Hasegawa, H., Hyodo, T., Ito, S., Asano, E., Yuang, H., Funasaka, K., Shimokata, K., Hasegawa, Y., Hamaguchi, M., et al. 2011 ARHGAP18, a GTPase-activating protein for RhoA, controls cell shape, spreading, and motility. Molecular Biology of the Cell. 22, 3840-3852. (10.1091/mbc.E11-04-0364)

26. Abouhamed, M., Grobe, K., San, I. V. L. C., Thelen, S., Honnert, U., Balda, M. S., Matter, K., Bahler, M. 2009 Myosin IXa Regulates Epithelial Differentiation and Its Deficiency Results in Hydrocephalus. Molecular Biology of the Cell. 20, 5074-5085. (10.1091/mbc.E09-04-0291)

27. Dai, B., Zhang, X., Shang, R., Wang, J., Yang, X., Zhang, H., Liu, Q., Wang, D., Wang, L., Dou, K. 2018 Blockade of ARHGAP11A reverses malignant progress via inactivating Rac1B in hepatocellular carcinoma. Cell Commun Signal. 16, 99. (10.1186/s12964-018-0312-4)

28. Howe, E. N., Cochrane, D. R., Richer, J. K. 2011 Targets of miR-200c mediate suppression of cell motility and anoikis resistance. Breast Cancer Res. 13, R45. (10.1186/bcr2867)

29. Abdi, H., Williams, L. J. 2010 Principal component analysis. Wiley Interdisciplinary Reviews: Computational Statistics. 2, 433-459. (10.1002/wics.101)

30. Milligan, G. W., Cooper, M. C. 1986 A Study of the Comparability of External Criteria for Hierarchical Cluster Analysis. Multivariate Behav Res. 21, 441-458. (10.1207/s15327906mbr2104_5)

31. Lawson, C. D., Ridley, A. J. 2018 Rho GTPase signaling complexes in cell migration and invasion. J Cell Biol. 217, 447-457. (10.1083/jcb.201612069)

32. Daugelaite, J., O’ Driscoll, A., Sleator, R. D. 2013 An Overview of Multiple Sequence Alignments and Cloud Computing in Bioinformatics. ISRN Biomathematics. 2013, 1-14. (10.1155/2013/615630)

33. Madeira, F., Park, Y. M., Lee, J., Buso, N., Gur, T., Madhusoodanan, N., Basutkar, P., Tivey, A. R. N., Potter, S. C., Finn, R. D., et al. 2019 The EMBL-EBI search and sequence analysis tools APIs in 2019. Nucleic Acids Res. 47, W636-W641. (10.1093/nar/gkz268)

